# Thrombin-preconditioned Mesenchymal Stem Cell-Derived Exosomes For Wound Healing in vitro and in vivo

**DOI:** 10.1101/2025.07.05.663313

**Authors:** Liu Yang, Minming Lou, Hongwei Wang, Shuo Zhang, Jie Ma

**Affiliations:** Institute of Biomedical Science, Tianjin Kang Ting Biological Engineering Co., Ltd., Tianjin 300385, P.R. China

**Keywords:** Mesenchymal stem cells, Umbilical cord derived MSCs (UCMSCs), Thrombin, Exosomes, Wound Healing, in vitro and in vivo

## Abstract

**Background and Objectives:** The skin is the largest organ of the human body, capable of protecting it from external harm. However, due to trauma, paralysis and other external factors, skin damage can occur, scars may form. Exosomes have regenerative functions and, as a cell-free therapy, show great potential for wound healing. In this study, we investigate that whether different umbilical cord mesenchymal stem cell exosomes (UCMSCs-Exos) can accelerate the healing of skin.

**Methods:** In our study, UCMSCs were cultured from newborns and Cultured with thrombin. Exosomes were isolated from mesenchymal stem cells culture supernatant by ultracentrifugation. The impact of exosomes on Hacat cell migration was evaluated in vitro. Additionally, the wound healing capacity of exosomes was evaluated in vivo using a mouse skin injury model.

**Results:** Compared to UCMSCs-Exos, T-UCMSCs-Exos significantly promoted cell proliferation and migration of cells. In vivo experiments demonstrated that T-UCMSCs-Exos can accelerate wound closure and enhance collagen maturation, promoting angiogenesis in the vascularized wound area. These results indicate that T-UCMSCs-Exos have a good potential for accelerating wound healing and minimizing scar formation.

**Conclusions:** Our research indicate that thrombin pre-UCMSCs, the production of exosomes significantly increased. The prepared T-UCMSCs-Exo can accelerate skin wound healing during the process of skin injury repair, promote angiogenesis, and facilitate the reconstruction of epidermis and dermis as well as hair follicle regeneration. These findings demonstrate that T-UCMSCs-Exos for skin wounds are a promising cell-free therapy that can be applied in the treatment of skin injuries.

## 1. Introduction

The skin is also the largest organ of the human body, forming its outermost layer. It covers the entire body and protects various tissues and organs from mechanical, pathogenic microbial, and chemical invasions. It has a protective effect on the body. It can also provide a protective function for the organism. The skin can sense external stimuli through neural transmission and analysis by the cerebral cortex. Not only is the skin a physical barrier to the body, but it is also an immune barrier, where skin immune responses are crucial in wounds and infections[1,2].

Skin also has the function of wound healing. When the body is subjected to external forces, the skin and skin tissue can regenerate. It requires the cooperation of a variety of cells and tissues, including cell proliferation, migration, matrix synthesis, skin contraction and growth factor related signaling processes[3,4]. The process of skin wound healing mainly includes hemostasis, inflammation, cell proliferation and tissue remodeling[5]. Most wounds heal within one to two weeks, but for certain chronic injuries that are difficult to heal, such as those in diabetic patients or those who have been bedridden for a long time, the process of healing can be prolonged. Chronic wounds suffer from prolonged healing difficulties, leading to scar formation. In chronic wounds, the remodeling of the extracellular matrix is delayed, angiogenesis decreases, and there may be some defects in the epithelium[6]. Severe or deep wounds that do not heal for a long time can lead to the regeneration of skin tissue. Now medicine is trying to find materials that can replace skin tissue, such as 3D printed skin, tissue engineering and advanced dressings[7,8].However, it is limited in terms of biocompatibility and cost, so how to solve the problem of wound healing has become an important issue in today’s research.Exosomes are extracellular vesicles with a diameter of 30-150 nm that carry a variety of bioactive molecules, including proteins, RNA (such as miRNA), lipids and cytokines[9,10]. These molecules facilitate cell proliferation, migration, and differentiation through intercellular signaling and communication, enabling the regulation of biological functions in target cells to achieve repair. They play a crucial role in skin injury repair and have been extensively studied experimentally in regenerative medicine[11,12].

In recent years, a large number of literature has reported that stem cells can treat the ability of chronic wound healing[13].There are also some literature indicating that stem cell exosome plays a decisive role in the treatment process[14].However, the specific mechanism is not yet clear, so it is urgently necessary to study the mechanisms affecting the secretion of mesenchymal stem cell exosomes and how to efficiently obtain mesenchymal stem cell exosomes for their application in skin injury repair. Currently, exosomes are mainly obtained directly from the supernatant of stem cell cultures using ultrafiltration methods[15,16]. However, the large amount of cell culture supernatant used and the limited acquisition of exosomes to some extent restrict the preparation and application of exosomes. Further exploration is still needed in the future, including establishing effective and feasible preparation methods as well as application approaches. Therefore, it is essential to find other efficient ways to obtain exosomes to overcome these limitations and understand their underlying causes.

In this experimental study, thrombin was used to stimulate mesenchymal stem cells to obtain cell culture supernatants. Exosomes were extracted from the supernatant using ultracentrifugation for the first time, preparing functional thrombin-exosomes and verifying that thrombin significantly increased exosome production by inducing the VEGF signaling pathway. Both types of exosomes were used in both in vitro cell models and in vivo models to promote wound healing. Our research results show that T-Exos can enhance cell migration and promote wound healing in damaged skin.

## 2. Materials and methods

### 2.1. Preparation of cell culture supernatant

Human umbilical cord MSCs (UC-MSCs) were derived from human umbilical cord (Kangting, China). After rapid thawing in a water bath at 37°C, 2×10^6^ cells per cryovial were seeded in a T175cm^2^ culture dish (Corning,NY, USA) and routinely resuspended in DMEM/F12(DF12,HyClone,USA), supplemented with10% fetal bovine serum(FBS,Hyclone,USA) and 1% penicillin/streptomycin (Guoyao, China). The cultures were maintained at 37 °C in 5 % CO_2_ and 95% humidity, and presented fibroblast-like morphology. The medium was changed every 2-3 days and MSC were subcultured when they reached 80-90% confluence. MSCs at passages 3-4 were used in the experiment.The MSCs cells were employed to produce exosomes.

When the UCMSCs density reached 60%-70%, the cells were rinsed three times with phosphate-buffered saline (PBS). The cells were cultured in serum-free DMEM/F12 alone as a control group or treated with 10U/ml Thrombin (sigma USA) in serum-free DMEM/F12. Then incubated for 72 hours prior to supernatant collectionfor to prepare umbilical cord mesenchymal stem cell exosomes (UCMSCs-Exos) and thrombin-treated umbilical cord mesenchymal stem cell exosomes (T-UCMSCs-Exos).

### 2.2. Isolation and characterization of exosomes

Exosomes were separated by ultracentrifugation[17]. After starving for 72h, the UCMSCs supernatant was collected in ultracentrifugation tubes (HITACHI, Japan). Initially, cell culture supernatant was subjected to centrifugation at 2000RPM for 20 min at 4 °C to eliminate cellular debris, and collected the supernatant. The resulting supernatant was centrifuged at 10000RPM for 30min at 4°C, and collected the supernatant. Then filtered the resulting supernatant once with 0.45μm filter membrane and 0.22μm filter membrane,and collected the supernatant. The resulting supernatant was centrifuged at 10000RPM for 70min at 4°C with the supernatant being discarded. PBS was used to resuspend the cells to harvest both UCMSCs-Exos and T-UCMSCs-Exos. The freshly isolated exosomes were preserved at -80°Cuntil needed.

### 2.3. Exosomes characterization

The exosomes were identified by the marker proteins CD9, CD63, Tsg101 Hsp70 or CD81 using western blotting, as well as by using a transmission electron microscope (TEM, Hitachi T-7800) to verify the exosome presence. The protein concentration of the exosomal fraction was quantified with the BCA protein assay kit following the manufacturer’s instructions (Pierce, USA). The nanoparticle size were measured by Zetaview(Particle Metrix, Germany).

### 2.4. Verification of the ability of exosomes to promote migration

In this study, scratch healing was used to evaluate the migratory ability of exosomes. Cell migration was assessed using a scratch wound healing assay. Hacat cells were initially inoculated into a 6-well corning culture plate with a density of 3×10^5^cells/well. When they reached 90% confluence, cells were cultured in serum-free high glucose DMEM for 24 hours.Then cells were treated with 10mg/ml Mitomycin for 2h before creating a wound by scraping the cell monolayer with a yellow tip. The width of the wound was measured at 0, 12, and 24h intervals, both in two kind of exosomes.

### 2.5. Wound healing animal model

This study selected male BALb/c mice, aged 42 to 55 days and weighing 18 to 22 grams.

Before modeling, they were administered a single intraperitoneal injection of 1% pentobarbital. A full-thickness skin lesion with a diameter of 15 millimeters was created on their backs. After the skin lesion model was established, mice with uniformly shaped and sized lesions were selected for the experiment. They were randomly divided into three groups: the exosome treatment group, the functional exosome treatment group, and the PBS control group, with six mice in each group.

The drug solution was applied dropwise to the wound site, applied locally once daily for 10 days.

During this period, the healing process of the wounds was monitored and recorded. At the end of the study, the mice were euthanized, and histological evaluations were performed on their wound samples, including hematoxylin and eosin staining, Masson trichromatic staining, and immunofluorescence staining.

### 2.6. Monitoring of wound healing

After treatment, routine observations are conducted daily until the wound heals; photographs of the wound are taken at 0, 2, 4, 6, 8, and 10d. A ruler (with each mark measuring 1 mm) is placed along the edge of the wound during photography, with the lens parallel to the wound surface. The wound area is calculated using Image J image analysis software. The wound area is measured using the elliptical formula:

Wound Area=half major axis length × half minor axis length × π.

Wound healing is considered complete when the wound is entirely covered by epithelial tissue or new skin. The wound healing rate is calculated as follows:

Wound Healing Rate = (Initial Wound Area-Current Day Measurement of Wound Area) / Initial Wound Area × 100%.

### 2.7. Histological observation

After 10 days of treatment, the mice were humanely euthanized and their wound sites were collected for histopathological examination. A 15mm²square piece of wound tissue was taken centered on the lesion, fixed in 4% formalin at room temperature for 24 hours, then gradually dehydrated and embedded in paraffin. The embedded samples were sectioned longitudinally with a thickness of 5 mm, and the sections were placed on polylysine-coated slides. HE staining was used to observe the structural characteristics of the skin at the wound site, granulation tissue, neovascularization, and inflammatory cell infiltration. Masson staining was used to examine collagen fiber deposition and arrangement at the wound site.

Immunofluorescence techniques were employed to detect the expression of CK14 keratin and Collagen I at the wound site. The prepared samples were observed and photographed under a confocal microscope, and further research analysis was conducted using Image J software.

### 2.8. Statistical analysis

All results are presented as mean±standard deviation and data analysis was performed using Graphpad Prism 8 software. Statistical significance was determined by Student’s t tests or one-way analysis of variance to evaluate the T-UCMSCs-Exo treatment effect.When the data did not meet normal distribution or histopathological scoring criteria, non-parametric rank sum tests with multiple comparisons among samples were used. Differences were considered statistically significant if P<0.05.

## 3. Results

### 3.1. Characterization of thrombin pre-MSCs

First, we evaluated the characteristics of thrombin-pre-treated stem cells(T-pre-MSCs) and UC-MSCs were isolated from human umbilical cord tissue and treated with 10 U/ml thrombin for 72 h.After thrombin treatment, UCMSCs exhibited a spindle-shaped morphology (Fig. 1A) and could differentiate into osteoblasts, adipocytes, and chondrocytes (Figure 1B). FACS analysis showed positivity for CD73, CD90, and CD105, while negativity for CD11a, CD34, and HLA-DR after thrombin treatment(Figure 1C). Annexin V FITC/PI staining showed that no significant apoptosis induction in MSCs upon exposure to thrombin (Figure 1D). These data confirm that thrombin pre-treated MSCs exhibit similar pattern in terms of both the extent and level of their original MSC characteristics.

**Fig. 1.**
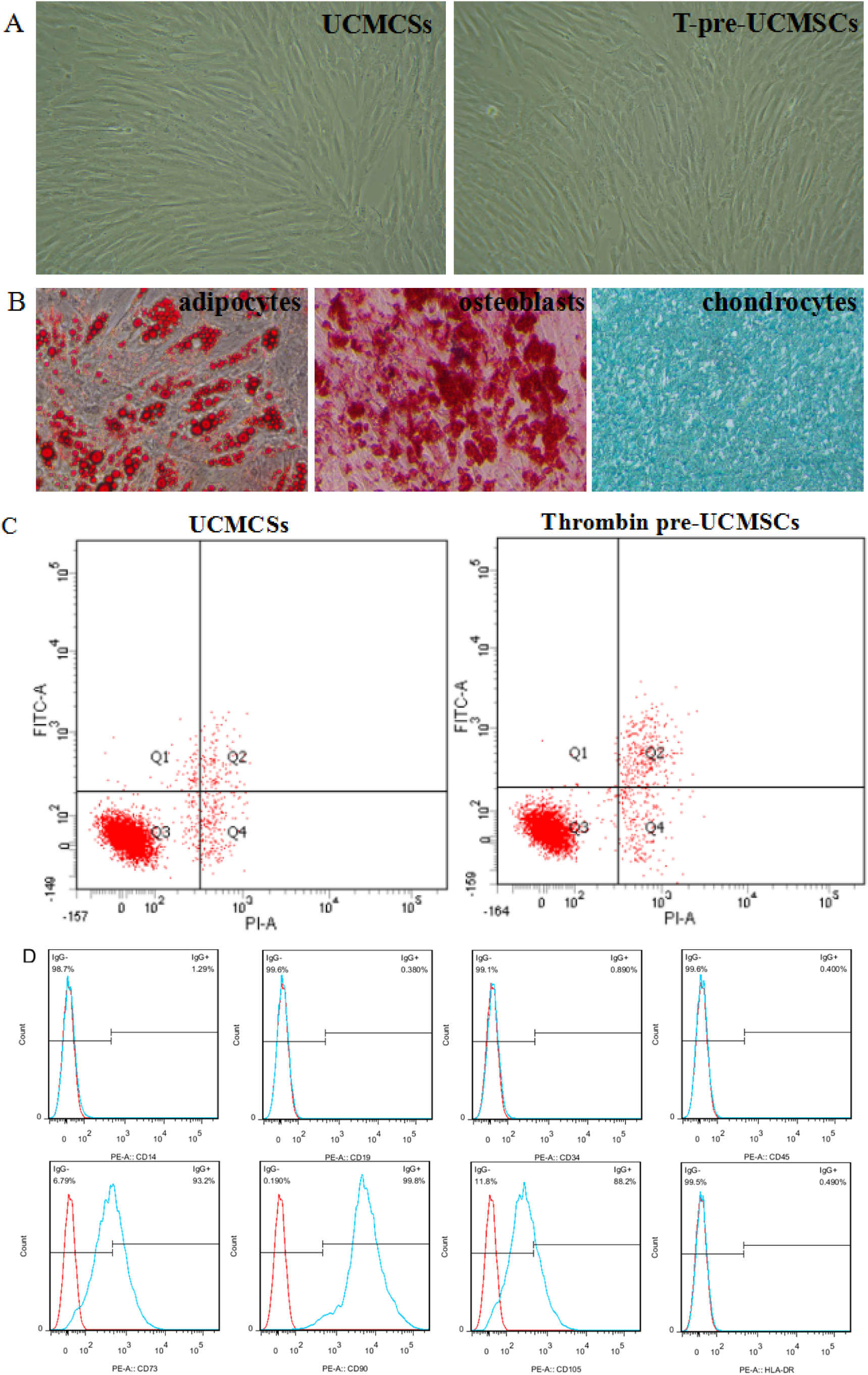
Characterations of Thrombin pre-MSCs. a Representative photomicrographs of UC-MSCs and Thromin pre-MSCs in culture displaying similar spindle shaped morphology. Scale bar 100 μm. B Thrombin pre-MSCs differentiation into adipocytes and chondrocytes. C After appropriate Thrombin stimulation, UC-MSCs apoptosis didn’t show clearly change by flow cytometry analysis of Annexin V and PE staining. D Immunophenotype of Thrombin pre-MSCs by flow cytometry. Data are presented as the mean ± SEM of three separate experiments.

### 3.2. Extracellular exsomes from hUCMCs culture supernatant exhibit exosomal traits

According to the significant role of exosomes in each ess of skin injury repair, we investigated the effects of T-UCMSCs-Exos and UCMSCs-Exos on wound regeneration both in vivo and in vitro. ucmsC with thrombin, collected the cell culture supernatant after starvation, and prepared thrombin-functional exosomes using ultracentrifugation. Exosomes size were detected by a nanoparticle size analyzer, ranging from 50 to 150 nanometers (2A and 2B). Exosomes morphology were characterized by scanning electron microscopy(TEM), which were showing as a teacup with relatively uniform size distribution (Figures 2C and 2D). Exosome markers were characterized using Western blotting (Figure 3E). According to BCA assay, the total protein concentration of UCMSCs-Exos was 159.22 ±26.34μg/500ml while the T-UCMSCs-Exos was 790.40±23.98μg/500ml(Fig. 3F). The results indicate that human umbilical cord mesenchymal stem cell-derived exosomes were successfully isolated in the experiment and can be used for further research.

**Fig. 2.**
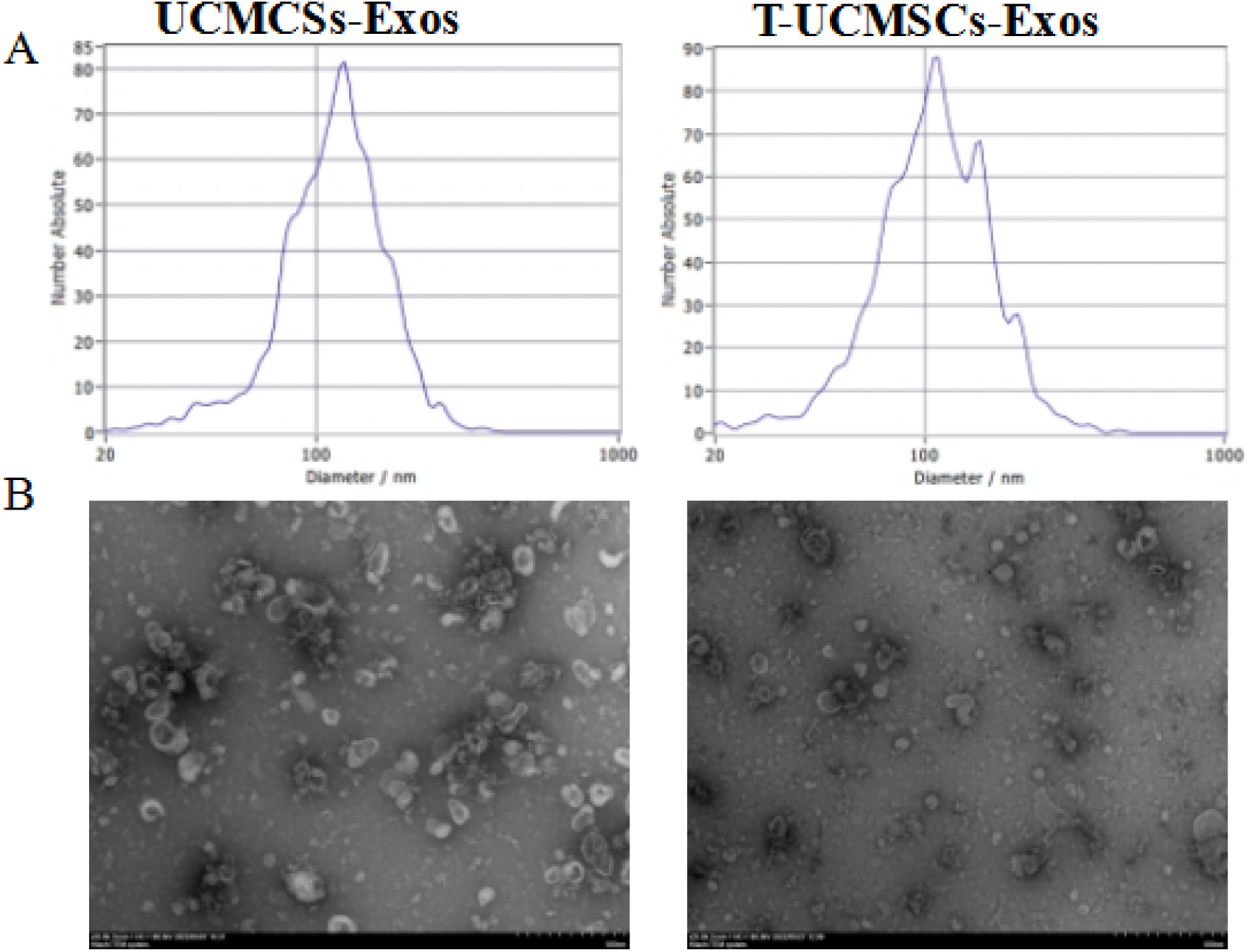
Nanoparticle tracking analysis (NTA) detection of exosome diameter. (A) Exosomes of exosomes observed under electron microscope.TEM showed that exosome exhibited a typical bilayer membrane structure (B)

**Fig. 3.**
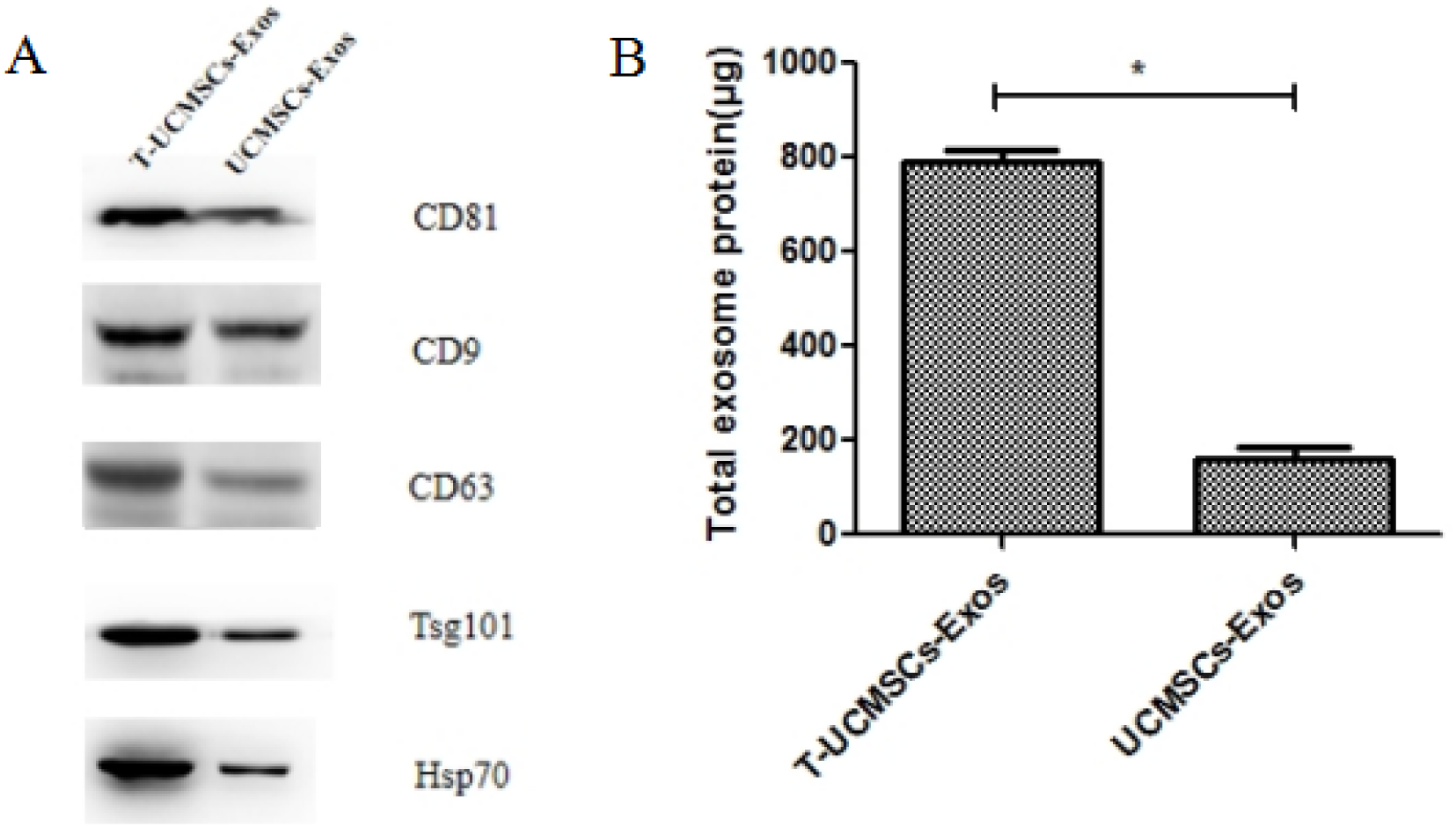
Western blot detection of CD81,CD9, CD63,Tsg101, HSP70 in exosomes(A); The total protein of the two kinds of exosomes was shown in Figure(B).

### 3.3. Human UCMSCs cell culture supernatant exosomes have an in vitro wound healing potential

To evaluate the effect of T-UCMSCs-Exos on human keratinocytes (Hacat) scratch assays were conducted. Hacat cells were seeded at a density of 2×10^6 cells per well in a 6-well dish. When the density reached 80%, a scratch was made using a 200μl pipette tip, and exosomes were added for treatment. The wounds were photographed at 6h and 24 h, and the wound closure percentage was calculated. The results showed that compared to the control group, thrombin-pre-treated exosomes significantly promoted the scratch healing of human keratinocyte-like cells Hacat (Fig.4)

**Fig. 4.**
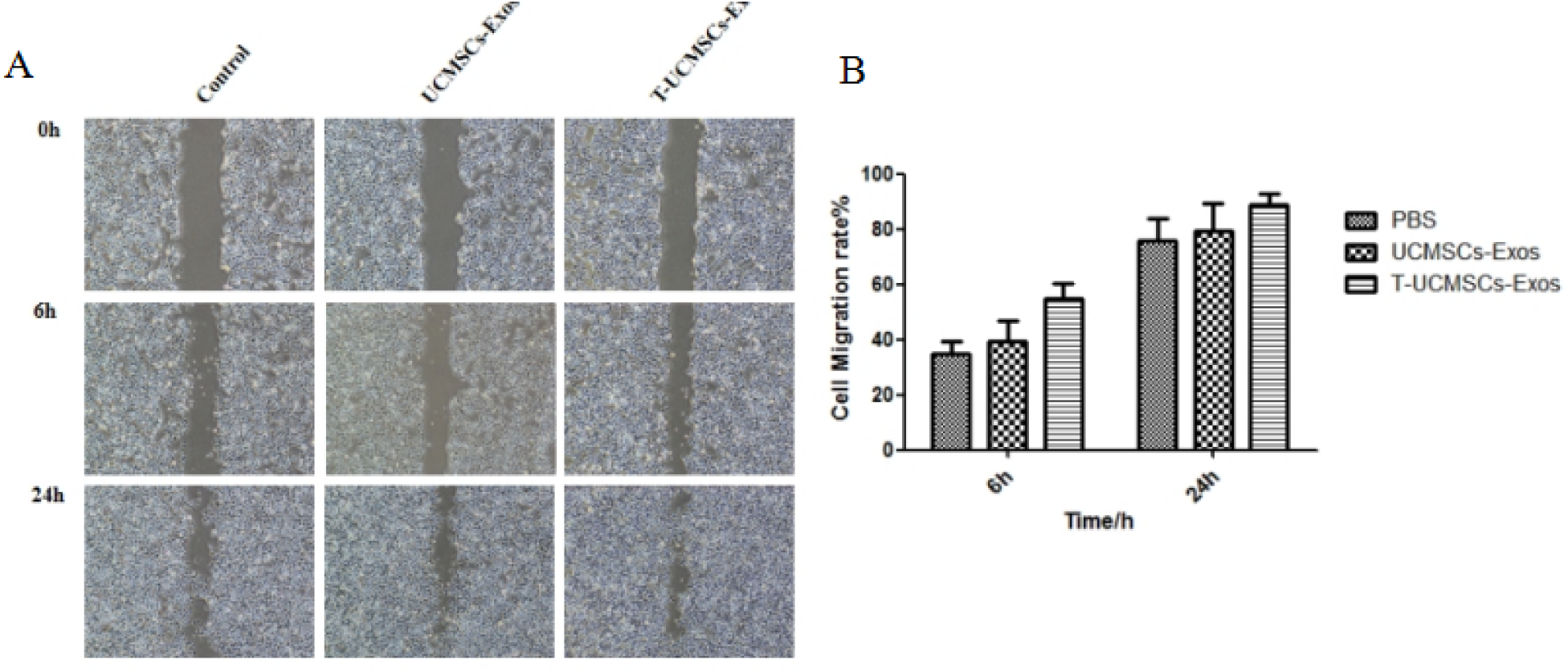
Evaluation of cell migration in vitro scratch assay. Photos of wound healing assay revealed that Hacat cells in the presence of exosomes displayed higher motility than nontreated cells as a negative control group after 24 h (upper panel, scale bar: 200mm). The lower panel shows the migration rate of cells that was assessed based on the distance of the selected wounded area at time intervals of 6h and 24 h and the percent of wound closure was determined for each time point. (n=, mean±SD). The cell-free zone was analyzed by image analysis software (Image J).

### 3.4 Human **UCMSCs** culture supernatant exosomes promote cutaneous wound healing in vivo

In the Balb/c mouse model, we investigated the role of exosomes. For this purpose, a circular full-thickness skin defect was created on the back skin of the mice (Figure 5). The mice were divided into three groups: PBS group, UCMSCs-Exos group (20 μg/100 μl), and T-UCMSCs-Exos group (20 μg/100μl), with no infection during treatment. At 8d post-treatment, the scabs in the T-UCMSCs-Exos group had fallen off, new epithelial tissue had formed, and the residual wound was minimal; the control group still covered with eschar and showed significant inflammatory response. By 10d post-treatment, the wounds in the T-UCMSCs-Exos group had largely healed, with skin color approaching normal and hair regrowth at the edges; the control group still had some eschar covering, but granulation tissue was growing in the wounds (Figure 5A). The average wound healing rate for the medium control group was (89.76 ± 3.55)%%, while that for the T-UCMSCs-Exos group was (97.62 ± 1.29)%%. The average wound healing rate of the UCMSCs-Exos group was higher than that of the control group (94.26 ± 1.47)%% (Figure 5B).

**Fig. 5.**
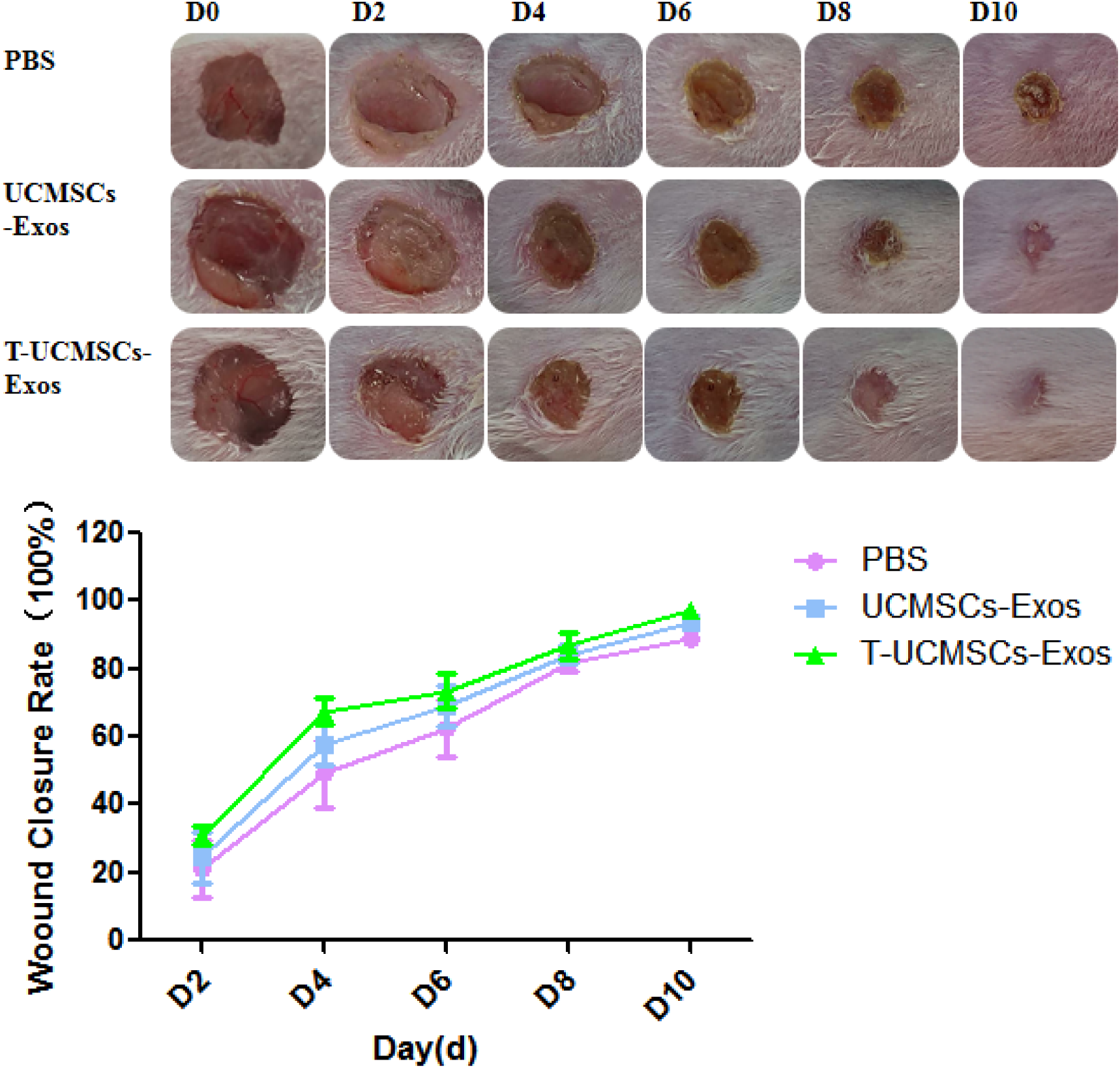
Showed that UCMSCs-Exos promoted wound healing. T-UCMSCs-Exos group exhibited the most significant improvement (A,B).

### 3.5. Human UCMSCs culture supernatant exosomespromote reorganization of wound tissue in vivo

The skin tissue samples obtained after treatment were stained with HE and Masson, and the length of wounds that were not fully healed was quantitatively analyzed using Image J. The results showed that the unhealed length in the PBS control group was 7.066±0.54mm, while the T-UCMSCs-Exos group was 2.425 ± 0.387mm, a decreasion of 65.68%. In the UCMSCs-Exos group the wound length were 4.896±0.892mm, a decreasion of 30.71%. The injection of exosomes accelerated wound healing (Fig.6A).

**Fig. 6.**
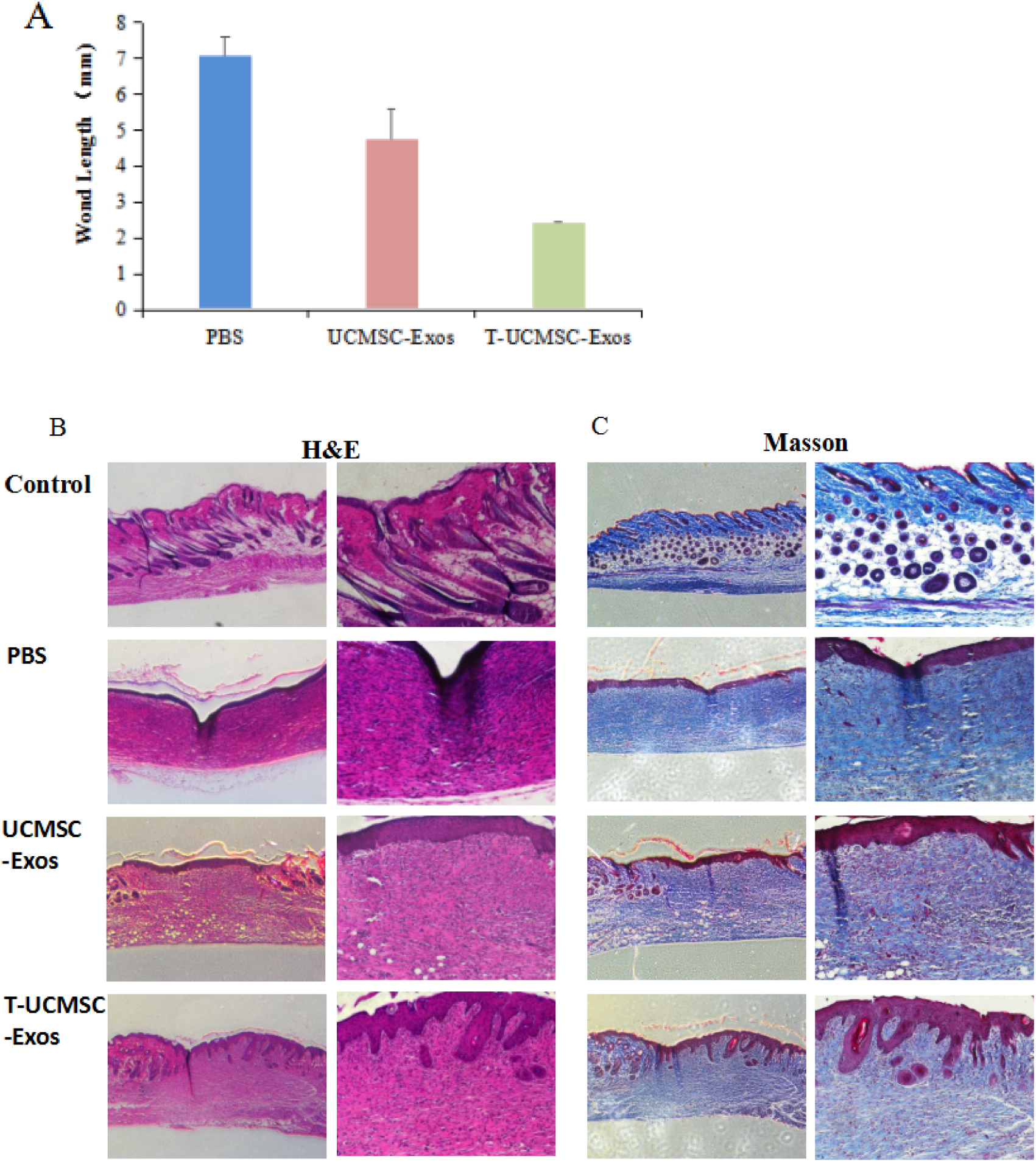
The mean length of wounds in each group of mice was measured(A) H&E and Masson’s staining were used to evaluate re-epithelialization and collagen remodeling (B,C).

H&E results showed that 10 days post-treatment, the epidermis was intact in all groups. In the T-UCMSCs-Exos treatment group, the skin layers were clearly defined and uniformly thick, with abundant collagen, hair follicles, and sebaceous glands in the dermis, showing normal morphology and no significant abnormalities. In the PBS model control group, which had a slower healing rate, the epidermis grew very thinly, with localized thickening of the epidermis, reduced destruction of hair follicles and sebaceous gland structures, disordered cell arrangement, and necrosis of muscle cells in the myofibrils, accompanied by minimal inflammatory cell infiltration (Fig. 6B).

Masson In the 10d after treatment in the dyeing, T-UCMSCs-Exos could promote the deposition and reorganization of collagen in the damaged wound, which was more loosely arranged than normal; the collagen in the PBS control group decreased significantly, with light coloring and uneven distribution, and the collagen fibers were crisscrossed and uneven (Fig. 6C).

The results showed that T-UCMSCs-Exos had the highest degree of fibrosis and epithelial thickness, which was related to the parallel transfer of a large number of growth factors to skin target cells by exosomes, which was beneficial to the healing of early wounds and enhanced the regeneration of skin tissue and collagen formation in full-thickness skin injury.

### 3.5 T-UCMSCs-Exos increased CK14 and Collagen I related proteins expression in wound healing

IHC study for the expression of CK14 and Collagen I(Col I) were performed to evaluate the extension of vascularization as shown in Fig. 7A.CK14 is rarely detected in the control group. In contrast, CD14 expression in the exosome group is significantly higher than in the model group, especially in the T-UCMSCs-Exos group. Meanwhile, the relative immunofluorescence values of CK14 in the control group is 1756.67±157.36, the PBS is 1499.60±262.43, the UCMSCs-Exos is 1940.92±350.61, and T-UCMSCs-Exos shows high CK14 expression at 2806.94±292.34 (p<0.001) (Fig.7B).The same results were obtained in immunofluorescence staining of Col I (Fig.7C).

**Fig. 7.**
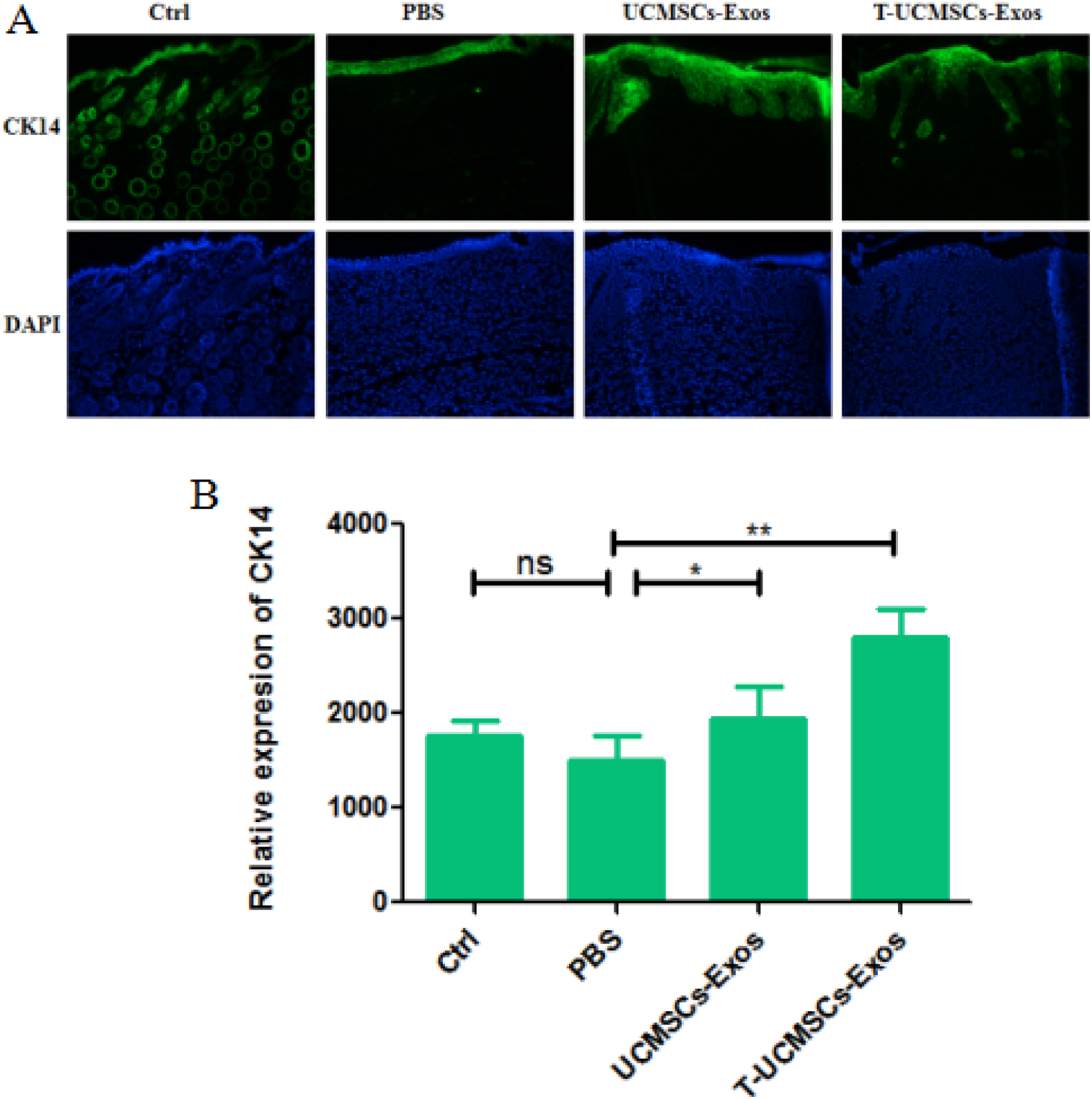
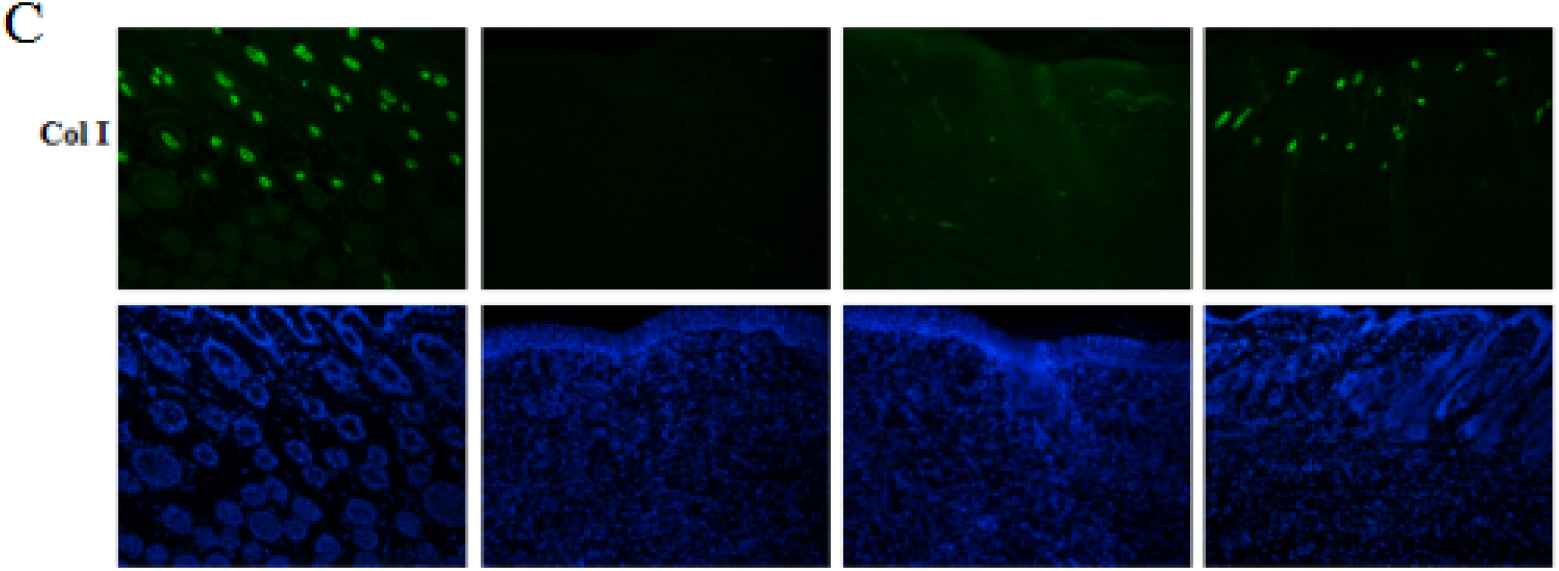
Histological analysis of the wounds in different experimental conditions on day 10. (A) CK14 staining of wounds from different groups (left panel). Quantitative evaluation of the percentage of CK14 positive area (right panel).(B) CK14 staining of wounds from different groups (left panel). Quantitative evaluation of the percentage of CK14 positive area (right panel). Skin tissue was obtained at day 10 post-surgery (left panel). (C) CK14 staining of wounds from different groups.Quantitative evaluation of the percentage of collagen deposition (right panel). *p < 0.05, **p < 0.01.

## 4. Discussion

The result of present research showed that human thrombin pre-UCMSCs cell culture supernatant exosomes have an significantly promoted the scratch wound closure of mouse human Hacat cells compared to the UCMSCs cell culture supernatant exosomes and control group. The results showed that the feature of thrombin pre-UCMSCs is similar to UCMSCs.Also in preclinical study, the effects of T-UCMSCs-Exos on wound healing study in healthy mouse, safety was assessed and no sign of inflammation and infection were observed.The efficacy of exosome application to the skin surface of mice was confirmed by reepithelialization, collagen fiber deposition, and granulation tissue formation after 10 weeks. More over, the rate of wound closure was higher compared to the control groups.

Angiogenesis in wound area has been shown by CK14 and collagen I antibody staining and confirmed percentage of healing was uncomprabale to control group. Significant reduction in wound size was detected after two weeks in both group.

The skin is the largest organ of the human body, covering the entire external surface and serving as the first line of defense for protecting the organism.The skin is very thin; when excluding subcutaneous fat tissue, the thickness of human skin ranges from about 0.5 to 4 mm[18,19]. Large wounds and chronic wound healing can lead to lifelong disabilities and other significant public health issues. Skin injuries mainly include burns, scalds, skin ulcers, UV radiation damage, diabetic wounds, eczema wounds, and tumor wounds[20]. Regardless of the type of tissue injury, it is a long and complex wound healing process that requires multiple tissues and cells in the body to work together to repair, regenerate, and rebuild damaged tissues, restoring their original functions[21–22]. And as paracrine substances in stem cell culture, exosomes have been found to possess advantages such as high differentiation potential, high survival rate after transplantation, and no significant adverse reactions compared to stem cells[23–25]. They contain a large number of proteins, cytokines, and bioactive substances, which play important roles in promoting the formation of new blood vessels and anti-inflammatory effects.Exosomes are extracellular vesicles secreted by various cells, with sizes ranging from 40 to 100 nm.Exosomes contain a large amount of proteins, lipids, transcription factors, as well as DNA, mRNA, and miRNA[26–27], which act on target cells by binding to receptors or fusing with the plasma membrane, thereby exerting their functions. Exosomes can transport various proteins, mRNA, and non-coding RNAs to regulate the activity of recipient cells. It plays an important role in the healing of skin wounds[28–30].

The healing process consists of three consecutive and overlapping stages: the inflammatory phase, the proliferative phase, and the remodeling phase. Stem cell-derived exosomes can promote cell proliferation and neovascularization, facilitate the function of fibroblasts, and enhance vascularization of endothelial cells, thereby aiding wound healing and promoting skin injury repair[31–33]. During the wound repair process, the transition from the inflammatory phase to the proliferative phase is characterized by the migration and proliferation of fibroblasts, keratinocytes, and endothelial cells, as well as angiogenesis, leading to granulation tissue formation and reepithelialization[34,35]. Exosomes can deliver growth factors and cytokines through this pathway. Studies have found that TGF-b and other molecules promote the proliferation, migration and formation of capillary endothelial cells, forming a network of capillaries, which is crucial for skin wound healing[36,37]. Research has found that mesenchymal stem cell exosomes contain abundant Wnt proteins, which can activate the β-catenin signaling pathway[38,39]. At the same time, by activating the Erk1/2 signaling pathway, they positively regulate the expression of angiogenic genes such as VEGFA, Cox-2, and FGF2, thereby promoting the regeneration of blood vessels, collagen synthesis, and skin injury repair in deep injuries of second-degree burn rat models[40].Exosomes extracted from the supernatant of mature umbilical cord mesenchymal cells through cell culture processes exhibit significant regenerative potential in stem cells[41,42]. However, clinical applications of exogenous stem cells face legal and ethical restrictions. Compared to cell transplantation, exosomes have many advantages, such as being stable and easy to preserve, non-immunogenic for allogeneic transplantation without rejection, and exosome-mediated cell-free therapy offers greater stability and long-term storability, with no risk of tumor formation and lower immunogenicity[43–45]. However, there are also some disadvantages that prevent exosomes from being widely recognized in preclinical studies, such as the low number of exosomes from mesenchymal stem cells, making it difficult to establish large-scale production systems, and the high usage volume of exosomes in clinical applications, which cannot meet clinical needs. Targeted research often requires engineering modifications, and the safety and efficacy of drug delivery need to be verified, along with the validation of delivery routes and so on.

In summary, our research explored a new method for preparing exosomes using thrombin to enhance their yield. This project established methods for preparing exosomes from stem cell supernatants and developed a quality control system. We studied the stability of exosomes in stem cell supernatants and investigated the mechanism by which thrombin promotes the secretion of exosomes from stem cells. By utilizing chemical substances in conjunction with cell culture media, we increased the yield of exosomes. Through effective evaluations of exosome treatments at both cellular and animal levels, we confirmed that treated thrombin-exposed exosomes have better potential for promoting wound healing and determined the optimal dosage of exosomes. This work validates the application value of exosomes and provides a theoretical foundation for future clinical applications. However, exosomes are still in the developmental stage, and many mechanisms require further investigation. Combining exosomes derived from stem cells with organic biomaterials can enhance the healing process, offering new insights into exosome research.

## 5. Conclusion

Our study demonstrated that the production of exosomes by traditional cell culture is very low in the preparation of exosomes. After using thrombin to induce umbilical cord mesenchymal stem cells, the production of exosomes was significantly increased as about 5 times. The prepared T-UCMSCs-Exos can accelerate skin wound healing, promote angiogenesis, and promote epidermis and dermis reconstruction as well as hair follicle regeneration during the process of skin wound healing.These findings demonstrate that T-UCMSCs-Exos is a promising cell-free therapy for skin wounds and can be applied to the treatment of skin injuries.

## Acknowledgements

This work was supported by the Technology Bureau of Xiqing District, Tianjin(x9zdzx-202007).

## Data availability statement

The original contributions presented in the study are included in the article/supplementary material, further inquiries can be directed to the corresponding author.

## Author contributions

Ma Jie contributed to conceptualization. Lou Minming and Ma Jie contributed to methodology.

Yang Liu contributed to the formal analysis and project administration,manuscript editing, writing-review & editing. Hongwei Wang contributed to the data curation, Investigation. Shuo Zhang contributed to the injury testing on the rats, tissue processing and mmunohistochemistry,validation and software. All authors read and approved the final manuscript.

## Publisher’s note

All claims expressed in this article are solely those of the authors and do not necessarily represent those of their affiliated organizations, or those of the publisher, the editors and the reviewers. Any product that may be evaluated in this article, or claim that may be made by its manufacturer, is not guaranteed or endorsed by the publisher.

## Ethics statement

This study was carried out in strict accordance with the recommendations in the Guide for the Care and Use of Laboratory Animals of the National Institutes of Health. The protocol was approved by the Committee on the Ethics of Animal Experiments of the Tianjin Kangting Biological Engineering Group Co. ltd(KIBS, KIBS-IACUC-2022-012-1).. All surgery was performed under sodium pentobarbital anesthesia, and all efforts were made to minimize suffering.

